# High resolution genetic mapping of causal regulatory interactions in the human genome

**DOI:** 10.1101/227389

**Authors:** Natsuhiko Kumasaka, Andrew Knights, Daniel Gaffney

**Affiliations:** Wellcome Trust Sanger Institute

## Abstract

Physical interaction of distal regulatory elements in three-dimensional space poses a significant challenge for studies of common disease, because noncoding risk variants may be substantial distances from the genes they regulate. Experimental methods to capture these interactions, such as chromosome conformation capture (CCC), usually cannot assign causal direction of effect between regulatory elements, an important component of disease fine-mapping. Here, we developed a statistical model that uses Mendelian Randomisation within a Bayesian hierarchical model framework, and applied it to a novel ATAC-seq data from 100 individuals mapping over 15,000 putatively causal interactions between distal regions of open chromatin. Strikingly, the majority (>60%) of interactions we detected were over distances of <20Kb, a range where CCC-based methods perform poorly. Because we can infer the direction of causal interactions, the model also significantly improves our ability to fine-map: when we applied it to an eQTL data set we reduced the number of variants in the 90% credible set size by half. We experimentally validate one of our associations using CRISPR engineering of the BLK/FAM167A locus, which is associated with risk for a range of autoimmune diseases and show that the causal variant is likely to be a non-coding insertion within a CTCF binding motif. Our study suggests that many regulatory variants will be challenging to map to their cognate genes using CCC-based techniques, but association genetics of chromatin state can provide a powerful complement to these approaches.

## Introduction

Three-dimensional (3D) interactions between regulatory elements are a fundamental process in gene regulation^1^. Understanding the guiding principles that control these interactions is a major research interest in genomics^2,3^. Long-range regulation also poses a significant challenge for studies of human disease because risk variants may be located many kilobases (Kb) from the genes they regulate, making causal variant identification difficult^4,5^. Chromosome conformation capture (CCC)-based techniques have enabled the generation of genome-scale maps of 3D contacts in human cells^6-8^. These maps have provided valuable insights into large-scale structure and organisation of chromosomes^9,10^, and often also provide useful information linking distal disease risk alleles with putatively regulated genes^11,12^. However, it can be hard to distinguish functional interactions, such as enhancer-promoter looping, detected using CCC-based methods from a background of random collisions^13^, which is particularly pronounced over distances of less than 20Kb^11^.

A complementary approach to mapping genome-wide 3D interactions is to utilise germline genetic variation. Quantitative trait loci (QTLs) mapping of chromatin traits can identify genetic variants that regulate chromatin both locally and distally, sometimes over distances of hundreds of kilobases^14-16^. These distal QTLs are known to be enriched in topologically associating domains^14,15^ (TADs), suggesting regulatory regions mapped by chromatin QTLs do indeed physically interact with each other. For fine-mapping of putatively causal variants identified in human disease studies, this approach has some attractive features. First, unlike CCC-based techniques, our ability to detect interactions between regulatory elements is not influenced by the distance between them. Second, QTLs identified in these studies can be naturally aligned with those from disease studies, using colocalisation^17^. Third, causal interactions between different regulatory elements can be deduced using Mendelian Randomisation (MR) technique^18-22^, where germline genetic variants can be used as an instrument to resolve relationships between different active regions. Here we develop and apply a Bayesian hierarchical model that incorporates techniques from MR to map causal regulatory interactions using a novel ATAC-seq data set from 100 unrelated individuals of British ancestry (Online Methods).

## Results

### The model

Our approach is based on previous observations that associations between genotype at the same genetic variant and chromatin accessibility often appear spread across multiple independent “peaks” of open chromatin, sometimes over great distances^16^. These signals can arise for multiple reasons (Fig 1A). One possibility is that two or more variants in tight linkage disequilibrium drive independent associations at different peaks (hereafter, referred to as “linkage”). Alternatively, a single variant might independently drive association signals at multiple peaks (referred to as “pleiotropy”). Finally, a single variant could modulate accessibility at one regulatory element that then alters accessibility elsewhere in the genome, an indication that these elements functionally interact in 3D space (referred to as “causality”). Our Bayesian approach classifies peak pairs within 500Kb of one another into hypotheses of linkage, pleiotropy, causality, a single QTL at either of the modelled peaks and a null hypothesis of no QTLs in either peak (Fig. 1A).

**Figure 1.**
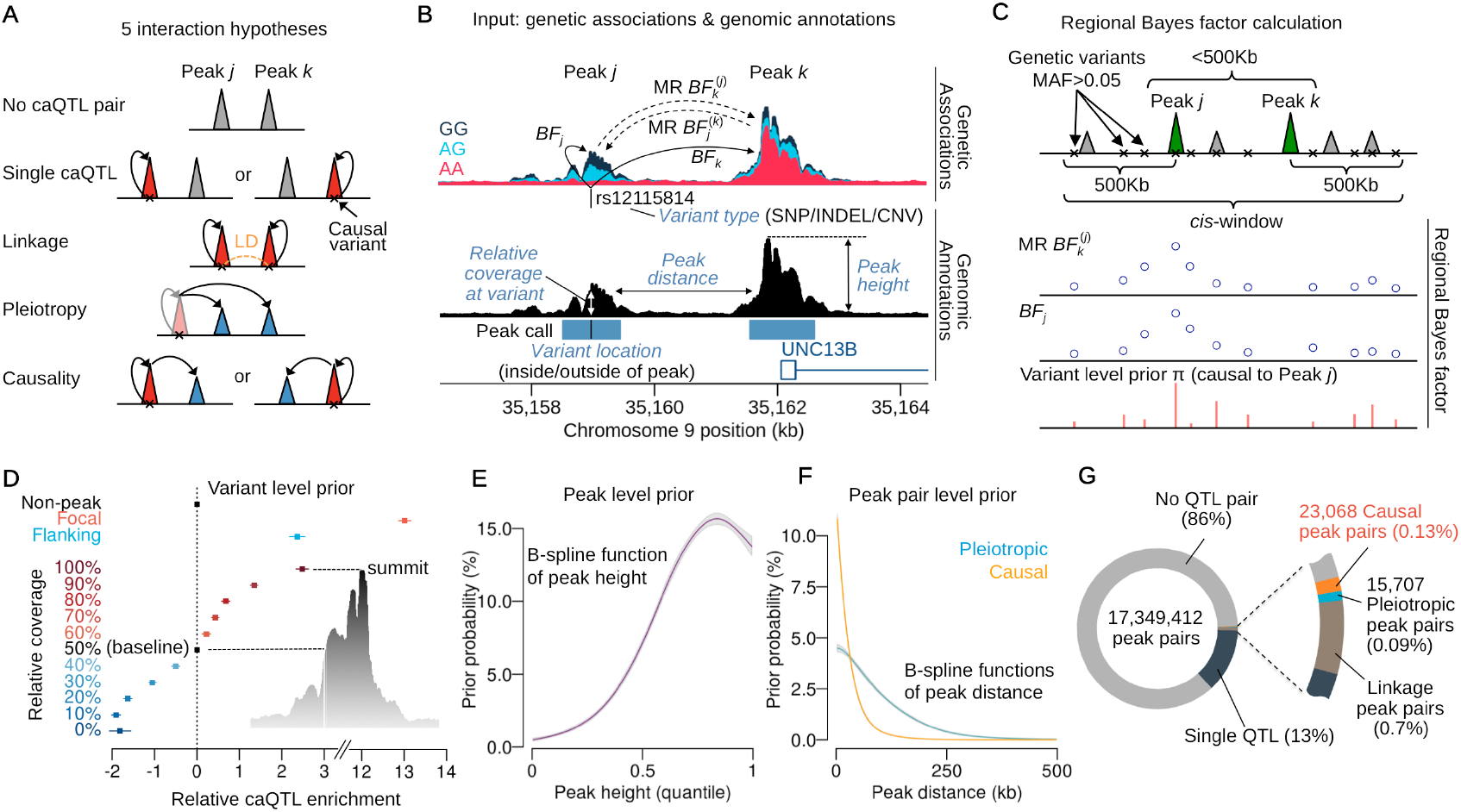
Overview of the pairwise hierarchical model and summary statistics. (**A**) The five main hypotheses of interaction between peaks. (**B**) Genetic associations with chromatin accessibility and related genomic annotations as input data. The Bayes factor (BF) of genetic associations with peaks (solid lines) are computed from the simple linear regression. The BFs of association between peaks (dashed lines) are computed using two stage least square method in the Mendelian Randomisation (MR). (**C**) For the *j-k* peak pair, BFs obtained in Fig. 1B are calculated for all variants in a *cis*-window and averaged as the regional Bayes factor (RBF). The schematic shows the two types of BFs across all variants were averaged by the variant level prior probability that the peak *j* is upstream of *k* (genetic variant is causal to peak *j*) to map causal interaction from *j* to *k*. (**D**) The estimated relative caQTL enrichments for genomic annotations used to compute the variant level prior probability in Fig 1C. (**E**) The estimated prior probability of a peak being a caQTL as a function of the peak height quantile among 277,128 peaks. The B-spline function was applied to capture non-linear relationship. (**F**) The estimated prior probability that a peak pair is pleiotropic or causal as a function of peak distance. Two different B-spline functions were applied. (**G**) The breakdown of mapped interactions according to Fig. 1A. The numbers are based on the sum of posterior probabilities.

There are three key features of the model. First, support for the hypothesis of a causal relationship between two peaks is computed using MR. Second, we use a hierarchical model^23^ in which prior probabilities depend on a range of genomic annotations at multiple model levels. Third, the model is empirical, such that the prior probabilities are learned as the likelihood is maximised across all peak pairs simultaneously. We model the relationship between genotype and a pair of chromatin accessibility peaks (Fig. 1B). To compute the pairwise likelihood for a given peak pair *j* and *k*, we calculate Bayes Factors (*BF_j_* and *BF_k_*) for the association between genotype at a putative causal genetic variant and chromatin accessibility at each member of the pair. For the hypothesis of causality we compute a Mendelian Randomisation Bayes Factors (MR 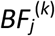 and MR 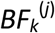) for the regression of chromatin accessibility in peak *j* on peak *k* (or vice versa) using the two stage least squares method^24^, with genotype at the given genetic variant as the instrumental variable (Fig. 1B). Using this information, the model can compare support for a causal effect of one peak on another, relative to the alternatives of pleiotropy, or two independent causal variants (linkage). Because the true causal variant is unknown, BFs are calculated for all variants in a *cis* window extending 500Kb 5’ and 3’ and marginalised by appropriate prior probabilities to derive the regional BF (RBF) (Fig. 1C). We use a “variant-level” prior probability of being a causal regulatory variant within *cis* window, based on variant location relative to and within a peak (Fig. 1D), and assuming a single causal variant in the window. We also model a “peak-level” prior on the probability of observing a caQTL, which is a function of peak height (Fig. 1E). This adjusts the support for QTL by the amount of data that supports that peak, an approach that is conceptually similar to independent hypothesis weighting^25^. Finally, we model a “peak-pair-level” prior that adjusts the support for pleiotropy or causality between two peaks, as a function of the distance between them (Fig. 1F) (see Online Methods for detail).

During initial model testing, we found that allowing for distance dependence between peaks provided a substantially better fit to the data than a uniform probability of interaction (Chi-square statistic=59,361 with *DF*=10; *P*<10^-12873^). Our data also strongly supported that causal interactions occurred over shorter ranges than pleiotropic (Fig. 1F). Following maximisation, the model outputs a posterior probability that a peak pair belongs to one of the interaction categories, including the posterior probabilities of a causal interaction (PPCs). For example, PPC_*jk*_ denotes the posterior probability that peak *j* regulates, or is “upstream” of peak *k*, while PPC_*kj*_ denotes the converse (*j* is “downstream” of *k*). In what follows, we use PPC to refer to the sum of PPC_*jk*_ and PPC_*kj*_.

### Mapped causal interactions

Of the 17 million peak pairs we considered genome-wide, 14% showed some evidence of genetic control, either a single QTL, linkage or some form of interaction (Fig. 1G). Summing over the posterior probabilities, we estimated that 23,036 peak pairs (0.13%) causally interact, of which 15,487 were high confidence, with posterior probability greater than 0.5. Our empirical prior suggested that the probability of any two peaks within 500Kb of each another interacting was 1.4% (Fig. 1F) suggesting that 1.23% or over 220,000 causal interactions remain to be discovered, although this is likely to be an underestimate. Following the initial round of interaction detection, we performed a *post-hoc* summarisation to identify directed acyclic graphs (DAGs) of causally interacting peaks in our high confidence call set (Online Methods). We identified 3,557 independent DAGs (Fig. S1A), of which 1,366 DAGs consist of more than 2 peaks up to 60 peaks as maximum at the MB21D2 locus (Fig. S1B) that we previously reported^16^. Fig. S1C is an example of DAG with three peaks in which the leftmost peak *A* regulates the flanking two peak *B* and *C* which are in pleiotropic relationship.

### The majority of causal interactions occur over sub 20Kb distances

Next we compared the 23,036 mapped causal interactions with loops inferred from Hi-C, promoter Capture Hi-C (Chi-C) and H3K27ac HiChIP applied to GM12878^10-12^ (Fig. 2A). More than 70% of the causal interactions we detected were between peaks located within the same topologically associating domains (TADs) called from Hi-C, an approximately 5-fold enrichment over genomic background (Figure 2B, C). Our interactions were also enriched for loops inferred from H3K27ac HiChIP and CHi-C data (7.7 and 1.4-fold, respectively), although the absolute numbers of overlaps were small (152 and 324, Fig 2C, B). This low overlap reflects the much shorter distances over which our interactions occurred (Fig. 2D): 63% were less than 20Kb distant from one another, compared with 7% of CHi-C interactions (Fig. 2D; Fig. S2A). Interaction distances were slightly longer when one member overlapped an annotated promoter (Fig. S2B; 49-54% < 20Kb). Thus, our approach revealed many functional three dimensional interactions are likely to be below the resolution of conventional CCC-based techniques. Our results also highlighted interactions that would be missed by promoter capture-based techniques. For example, there were 2,208 causal interactions between enhancers linked to the same baited promoter (Fig. 2B, C), and a further 561 causal interactions between two peaks located within the same CHi-C bait region. One example of a candidate short-range causal interaction at the promoter region of the MAP1B gene is shown in Fig. 2E. Here, a strong caQTL harbouring a putative causal SNP (rs1217817) is located immediately distal to the promoter flanking region that is causally interacting with the promoter peak with high confidence (PPC=1.0). Although this region overlaps a contact domain called from CHi-C, this interaction does not have strong statistical support (CHICAGO score 1.87) due to the short distance (<13Kb).

**Figure 2.**
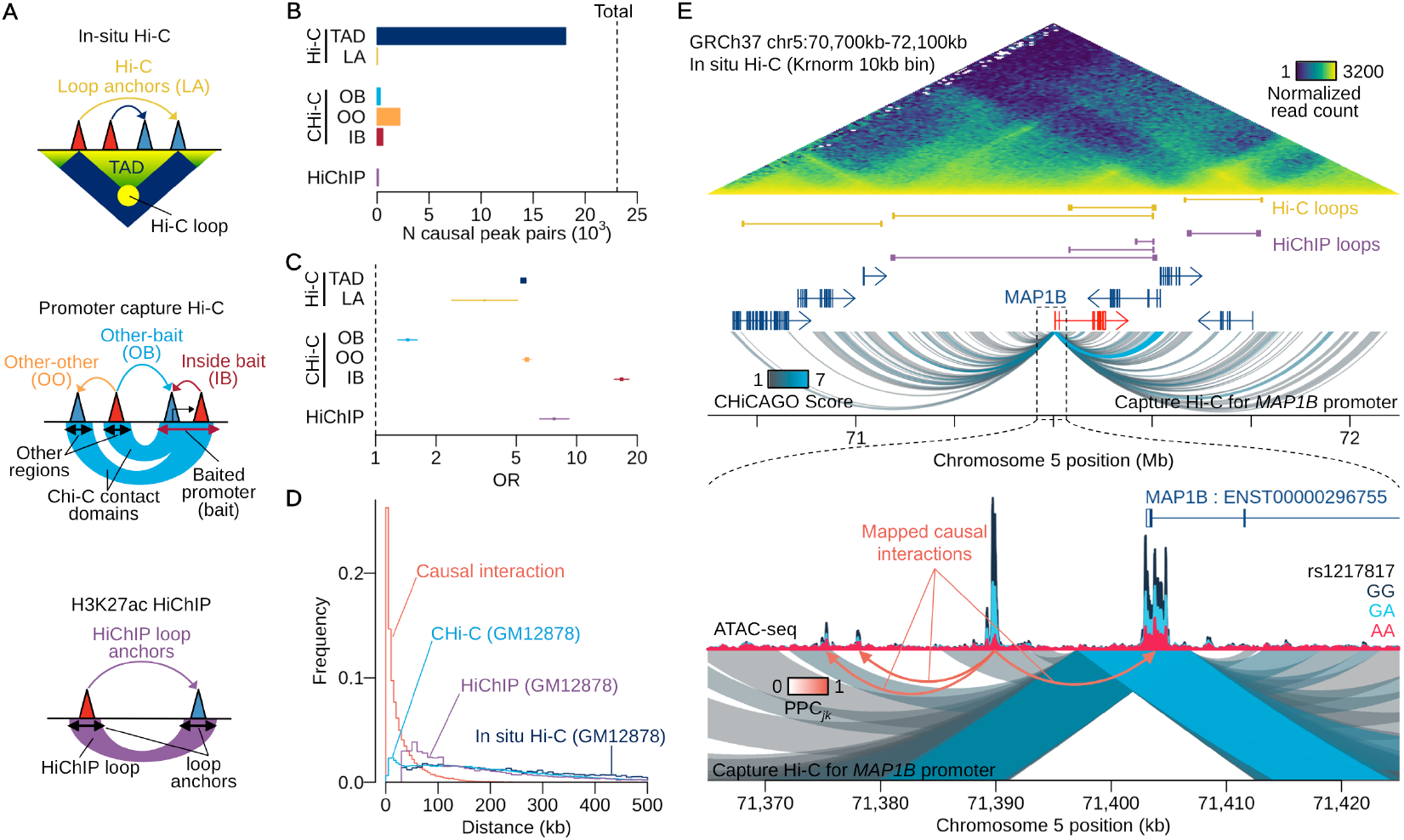
(**A**) Schematics of Hi-C, CH¡-C and HiChIP annotations. (**B**) The numbers of causal peak pairs overlapping with the annotations in Fig. 3A. Topologically associating domains (TAD), loop anchors (LA), other-other (OO), other-bait (OB), inside bait (IB). (**C**) Enrichment Odds Ratio (OR) of causal interactions with annotations. (**D**) Distribution of interaction length. For our interactions, we computed the distance between all 17 million peak pairs considered, and weighted these by the posterior probability of causality (PPC). (**E**) An example of causal interactions found in the promoter flanking region of MAP1B gene. There is a caQTL peak (with the QTL SNP rs1217817) 10Kb upstream of MAP1B promoter affects multiple open chromatin peaks including the promoter peak.

### Enhancer-enhancer and promoter-enhancer interactions are common

We next examined the functional classes to which the members of causally interacting regulatory elements belonged, using the ENCODE genome segmentation annotations for LCLs^26,27^ (Online Methods). The most frequent class of interactions (5,061 peak pairs, 22% of all interactions) were strong enhancers that appeared to regulate other element types, including other strong enhancers (1,531 peak pairs, 6.6%), a 2.5-fold enrichment (Fig. 3A, B). When we focussed only on variants that also altered gene expression, using 4,670 interacting peak pairs that jointly colocalised with an eQTL from the GEUVADIS data set (Online Methods), we found these were enriched (2.4-fold, *P*=6.4×10^-19^) for strong enhancer to active promoter interactions (Fig 3C, D). However, expression-associated variants were also enriched for interactions from active promoters to strong enhancers (2.2-fold) or between pairs of strong enhancers (2.2-fold enrichment) (Fig. 3D). One hypothesis is that many of these are mediated by transcriptionally induced changes in chromatin accessibility over the gene body, which create apparent interactions between a single upstream functional element and chromatin peaks throughout the transcribed region. A striking example of this potential phenomenon is found at the MB21D2 locus (Fig. S1C). This hypothesis is consistent with the observation that chromatin accessibility over the gene body is highly correlated with gene expression level (Fig. S3A. Furthermore, we found that peaks downstream of an active promoter were significantly enriched in the gene body (2.3-fold enrichment, *P* = 8.1×10^−24^; Fig. S3B; Online Methods) compared with peaks to the 5’ of the promoter.

**Figure 3.**
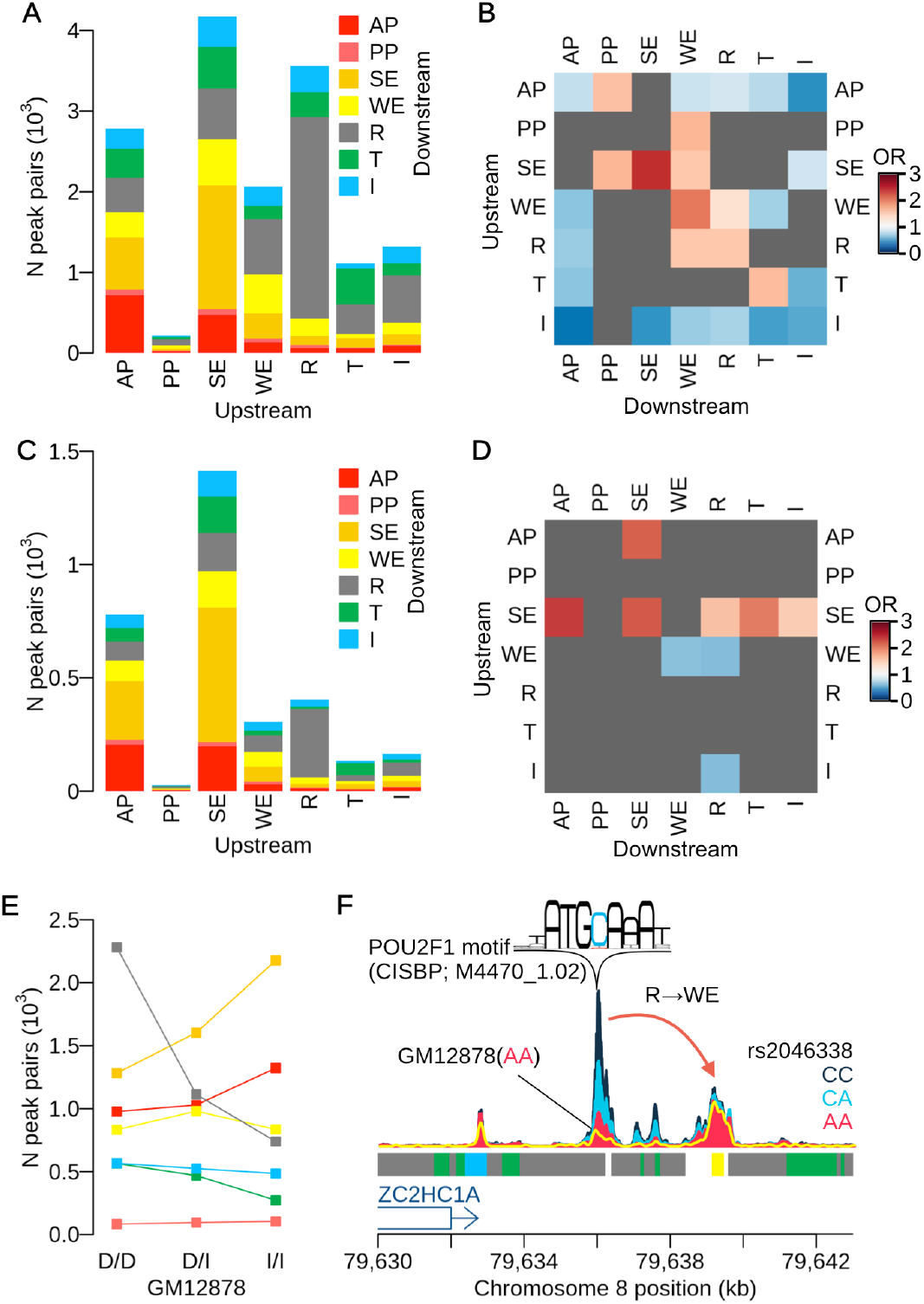
(**A**) The numbers of causal peak pairs overlapping ENCODE genome segmentation. Numbers of interactions were computed weighting by PPC_*jk*_. The ATAC-seq peaks are classified by 7 different regulatory categories: active promoter (AP); poised promoter (PP); strong enhancer (SE); weak enhancer (WE); repressor (R); transcribed region (T); and insulator (I). Each bar indicates upstream peak category and the colour code indicates downstream peak category. (**B**) Enrichment of ENCODE segmentation category pairs for our causal interaction. Heatmap shows the odds ratios for all combinations of segmentation categories at upstream and downstream peaks. The segmentation category pairs that were above FDR 10% or supported by less than 10 causal peak pairs are masked by gray. (**C**) The numbers of causal peak pairs that are jointly colocalised with one or more eQTLs overlapped with the ENCODE segmentation. (**D**) Enrichment of ENCODE segmentation category pairs for our causal interactions that are jointly colocalised with one or more eQTLs. (**E**) The number of peak pairs whose upstream peak overlaps with one of the seven segmentation categories, stratified by the genotypes of GM12878 at lead QTL variant (Online Methods). Each genotype is labelled as a combination of decreasing “D” and increasing “I” alleles according to the sign of QTL signal at the lead variant. Colour code is same as in Fig. 3A. (**F**) An example of causal interaction from a repressed region to a weak enhancer. The normalised ATAC-seq coverage is stratified by three genotype groups at rs2046338. The yellow line shows ATAC-seq coverage of GM12878 whose genotype is AA (decreasing homozygote) at rs2046338.

### Genetically-driven changes in the reference epigenome

We found a surprisingly large number of interactions (4,134 peak pairs) originating from within repressed regions (Fig. 3A). Preliminary analysis suggested that these might arise due to genotype effects on the reference epigenome annotation derived from a single individual (GM12878). To test this, we stratified all upstream peaks in causally interacting pairs based on whether their lead caQTL genotype in GM12878 was increasing homozygote, decreasing homozygote or heterozygote (Online Methods). Upstream repressed regions were highly enriched (3.1-fold) for decreasing homozygotes compared with increasing homozygotes (Fig. 3E), suggesting that in these cases a strong caQTL almost completely removes a region of open chromatin in GM12878, an example of which is shown in Fig. 3F.

### Causal interactions improve fine-mapping

Next we examined whether the information on causal direction of variant effects could be used to improve fine-mapping accuracy, using gene expression as a model quantitative trait. For each peak within a 1Mb *cis*-window around a gene TSS, we first computed the probability of master regulator (PMR) for each peak (see Online Methods). We then used a hierarchical model^23^ to compute the posterior probabilities of association (PPA) for eQTL variants with PMR and the following four other annotations: (1) inside or outside an ATAC peak, (2) eQTL variant location, relative to an ATAC peak coverage, (3) promoter CHi-C contacts, (4) HiChIP loops from promoter regions (see Online Methods for details). Genome-wide, the best performing annotation was the combination of PMR with ATAC peak status and variant location, which reduced the 90% credible set of eQTL variants by 65%, from 17 to 6 variants on average, compared with 11 variants for CHi-C, 10 for ATAC peaks and 8 for Chi-C combined with ATAC peaks (Fig. 4A). We examined the effect of adding information on the causal direction via the PMR, by comparing to variants annotated using ATAC peak data alone. The PMR annotation significantly reduced the credible set size (*P*<10^-49^, paired t-test). We then compared our results with data from massively parallel reporter assay (MPRA) performed in LCLs^28^. Here, we selected the lead eQTL variants, ranked by the eQTL PPAs for each annotation and asked how many overlapped a validated expression-modulating variant (an “emVar”) from the MPRA (Online Methods). We found the highest overlap (21.6% or 182 emVars) for the combined PMR, ATAC peak and variant location annotations (Fig. 4B). We show an example of this approach applied to a challenging locus in Fig. 4C, where a strong eQTL for the GPATCH2L gene is associated with more than 100 candidate regulatory variants in almost perfect LD. With no annotation information, the 90% credible set size at this locus is large, at 65 variants. Although different annotations produce varying effects, our model proposes a single SNP, rs74067641, as the likely causal regulator with the highest posterior probability (PPA=0.42). This is because this variant is located within what our model predicts to be a master regulatory peak located furthest upstream in the regulatory cascade (Fig. S4).

**Figure 4.**
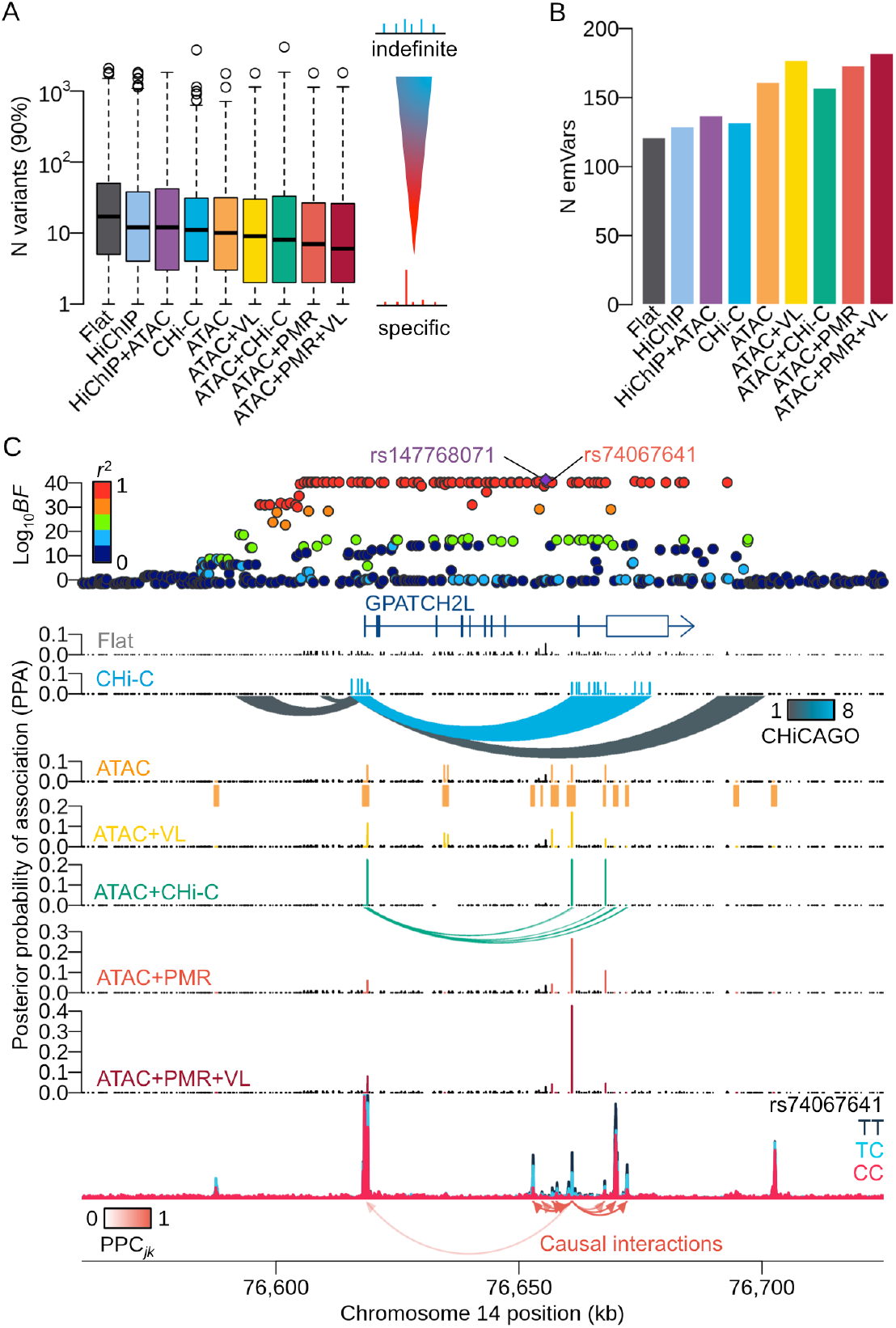
(**A**) Distribution of the number of variants in the 90% credible set, across all protein coding genes with more than one colocalised ATAC peaks (*N* genes=1,207) over nine different annotation combinations. FLAT: non-informative prior. ATAC: inside/outside an ATAC peak. HiChIP: HiChIP anchor regions. CHi-C: CHi-C contact domains. VL: variant location. PMR: probability of master regulator. (**B**) The number of expression-modulating variants (emVars) overlapping lead eQTL variants detected by the eQTL hierarchical model with various annotations. (**C**) An example of fine-mapped region with more than hundred of significant variants in almost perfect LD. The top panel shows negative Log_10_ Bayes factors of eQTL for GPATCH2L gene using gEUVADIS RNA-seq data. Each point is coloured by the degree of LD index (*r*^2^ value) with the index variant (rs147768071). The SNP (rs74067641) in the master regulatory peak shows the highest PPA with ATAC+PMR+VL annotation.

### CRISPR validation of a putatively causal variant at the BLK locus

Finally, we applied our method in an attempt to fine-map a challenging GWAS locus with contradictory evidence for multiple causal variants in previous studies. The BLK/FAM167A locus on 8p21 has a strong eQTL (gEUVADIS *P*<10^-26^ and 10^-46^ for BLK and FAM167A genes, respectively) in LCLs (Fig. 5A) that colocalises well with genome wide significant associations for systemic lupus erythematosus (SLE) and rheumatoid arthritis (RA) (Fig. S5.1A-B). Previous attempts to fine-map this locus have been hampered by multiple genetic variants in tight LD (Fig. S5.1C-D). Two SNP variants, rs1382568 and rs922483, located near the promoter of BLK gene, have previously been reported as putative causal variants of SLE that alter BLK expression in various B and T cell lines^29^. However, MPRA studies have pinpointed an alternative deletion variant (rs5889371) that might also potentially alter BLK expression in LCLs^28^. Nonetheless, two of the previously reported variants (rs5889371, rs1382568) are located in regions of low chromatin accessibility (Fig. S5E-G) and therefore less likely to causally influence BLK expression.

**Figure 5.**
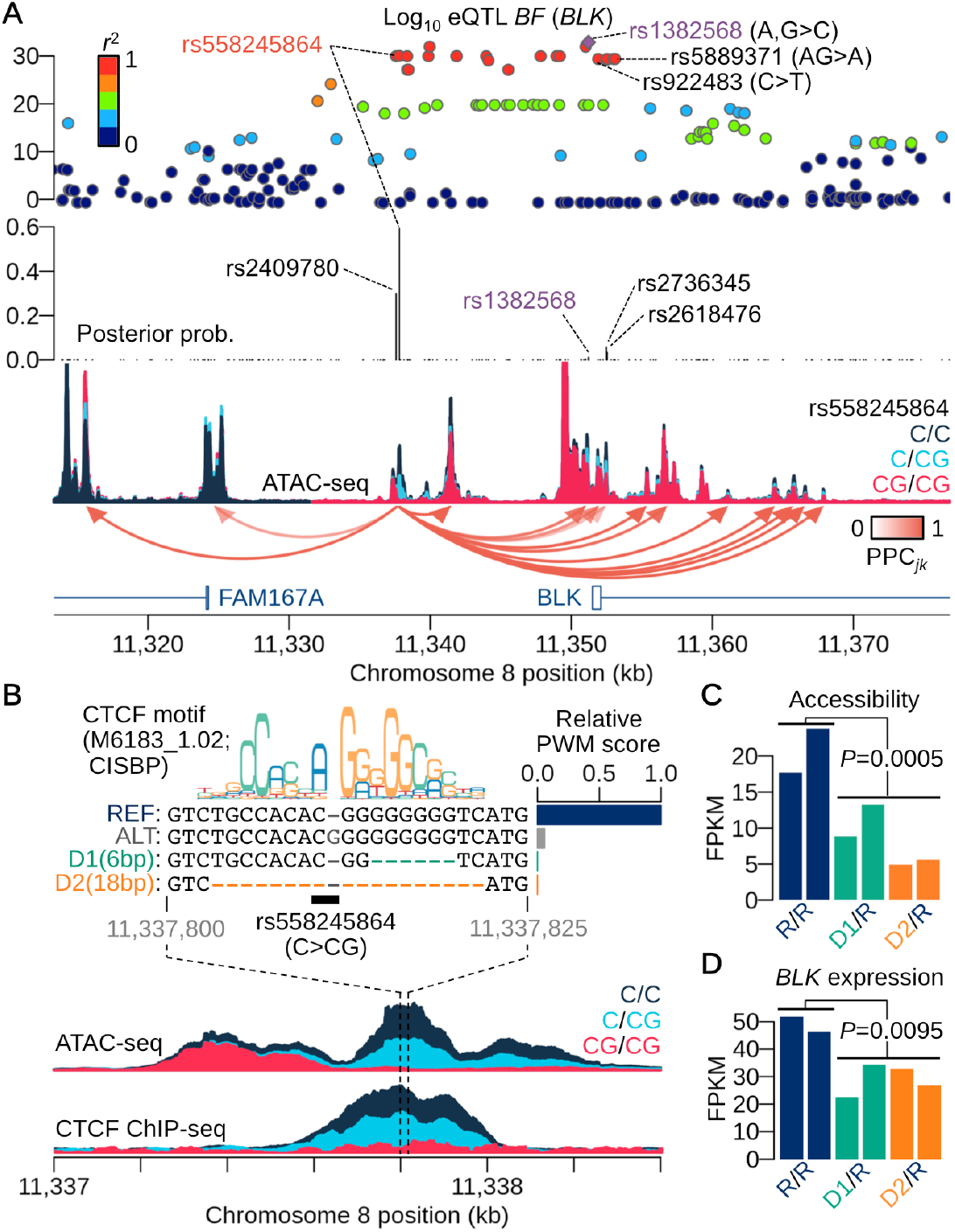
(**A**) A chromatin accessibility altering variant at the BLK/FAM167A locus. The top panel shows negative log_10_ Bayes factors of eQTL mapping for BLK gene using gEUVADIS RNA-seq data. Each point is coloured by the degree of LD index (*r*^2^ value) with the index variant (rs1382568). The middle panel shows the posterior probability of association (PPA) obtained from the full annotation model (Online Methods) in which the insertion variant rs558245864 shows the highest PPA. The bottom panel shows ATAC-seq coverage depth stratified by the insertion variant (rs558245864) with the causal interactions from the peak in which the insertion variant exists. (**B**) CRISPR engineered locus around the insertion variant. The insertion variant disrupts the CTCF binding site and attenuates the binding affinity (bar plot) calculated from the canonical CTCF binding motif (CISBP: M6183_1.02). Independent analysis of CTCF ChIP-seq binding QTL supports the result. CRISPR engineering was performed to generate two different deletions (D1 and D2) from the parental line (HG00142) whose genotype is reference homozygote at the insertion variant. The maximum CTCF binding affinity around the region after extracting the deleted sequences is lower than that of the alternative allele. (**C**) FPKMs at the focal peak for the two heterozygous deletion lines (D1: green and D2: orange) compared with the parental line with reference homozygote (R/R: navy). All lines were replicated twice. (**D**) FPKMs of BLK gene expression for the same lines in Fig. 5D.

We detected a single base pair insertion variant (rs558245864) located in a strong caQTL peak 14Kb upstream of the BLK promoter that interacted with 15 flanking peaks including several promoter peaks (Fig. 5A). The insertion variant showed the highest posterior probability (PPA=0.59) of any putatively causal eQTL variant for BLK gene (Fig. 5A). This variant is located at the middle of a canonical CTCF binding motif, with an extra “G” nucleotide decreasing the predicted CTCF binding affinity to almost to background (Fig. 5B). The direction of binding affinity change was consistent with the caQTL signal. This variant was also a CTCF ChIP-seq QTL (Fig. 5B), with 99.7% the probability of colocalisation between CTCF binding and the caQTL for this peak (Online Methods). We used CRISPR/Cas9 genome engineering to generate two different heterozygous deletion lines from a parental line that was homozygous for the high CTCF binding allele (Online Methods). These deletions overlapped the CTCF binding site: the 6bp deletion disrupts the right hand side of the binding site and the 18bp deletion that removes almost the entire motif (Fig. 5B). ATAC-seq and RNA-seq in the deletion lines revealed a significant down-regulation of chromatin accessibility at the focal peak compared with the parental line (*P*=0.0005) (Fig. 5C), and a concomitant down-regulation of BLK expression (*P*=0.0095) (Fig. 5D). We also observed decreases in accessibility at some neighbouring peaks (Fig. S5H) and increase in expression at the nearby FAM167A gene (Fig. S5I) as the eQTL results would have predicted, although it is not significant (*P*=0.18).

## Discussion

We have presented a novel approach to detect interactions between regulatory elements that utilises principles of Mendelian Randomisation technique within a Bayesian hierarchical model. We show that the majority of causal interactions within 500Kb occur over short distances (<20Kb), typically a region of low sensitivity for CCC-based techniques. Many of the interactions we detect are between enhancers, which we assemble into hierarchies of interacting regulatory elements. We demonstrate that our model can be used to identify hierarchies of regulatory elements within a region and prioritise putatively causal variants, validating a single locus using CRISPR/Cas9 editing.

The low frequency of long-range interactions we observed agrees with previous estimates from eQTL studies^23,30,31^. One question is why, given that most regulatory interactions detected using CCC-based methods over distances of 100Kb and above (Fig. 2D), large numbers of genetic variants operating at these distances have not been detected by QTL studies. This partly reflects the fact that QTL studies often test variants in a *cis* window of less than 1Mb^30-32^. However, even technical reasons seem insufficient as an explanation, given that the number of eQTL associations detected decreases dramatically by approximately 20Kb distant from the gene TSS^23,31^. One explanation is that there may be an underlying relationship between interaction distance and cellular frequency, such that long-range interactions occur in a relatively small number of cells in the population^30^. An observation supporting this hypothesis is the strong negative correlation between read coverage and distance in Chi-C data (Fig. S6A). It seems plausible that CCC-based methods could be more sensitive to rare, long-range regulatory interactions while variants residing in these elements have relatively weak effects^31^, requiring large sample sizes to detect when averaged across the entire cell population. If this hypothesis holds, our results highlight the importance of better characterisation of the mode of action of disease associated variants at the single cell level.

One of the limitations of our method is that regulatory elements lacking a common genetic variant that perturbs their function will be missed by our method. Additionally, interactions between genotype and regulatory elements further downstream appear to become harder to detect, perhaps due to additional biological noise. One example of this is the systematically lower genetic effect sizes (14 % decreasing) we found at downstream promoters (Fig. S6B, C).

Our study also revealed the genomic architecture of causal interactions between regulatory elements. In particular, we detected frequent interactions between annotated enhancer elements, many of which we hypothesise are mediated by an intermediate eQTL that alters chromatin accessibility globally across the gene body. Nonetheless, the enrichment of these interactions in gene bodies was modest, and we also found many examples of interactions that were not colocalised with eQTLs, and were located far from annotated genes (an example is shown in Fig. S6D). In a small number of cases (18 DAGs) we also found strong evidence (PPC > 0.5 for each enhancer pairs) that these occurred between multiple enhancers upstream of a promoter (*i.e*., SE➜SE➜AP). It is possible that some of these represent enhancer “seeding” events, where individual enhancers drive progressive activation of additional nearby elements^33^.

The approach we have developed allows for a natural prioritisation of variants in disease-associated loci. Although overlapping of disease associated variants with open chromatin can reduce credible sets, this frequently leaves many loci with tens of variants to characterise by direct experimental follow up. Assignment of the direction of effect between different peaks allowed us to identify smaller sets of plausible candidate variants by identifying “master regulatory” regions. Although here we have focused on ATAC-seq data, we believe our model can be readily extended to other types of chromatin-based assay, in particular ChIP-seq for histone modifications^14,15^. Some limitations of this approach might include a greater difficulty in assigning causal variants based on their location within a ChIP-seq peak, which will typically be in a nucleosome depleted and therefore low read coverage^14^ (Fig. S6B). However, we anticipate that, applied to existing data sets from primary cells, such as that generated by the BLUEPRINT initiative^34^, that our approach will be a valuable tool in dissecting the molecular architecture of specific GWAS loci.

## Online methods

### ATAC-seq in LCLs

We collected 76 LCL samples of British ancestry (1000 Genomes Project GBR) that we combined with 24 LCL samples previously sequenced^16^. We also performed an additional ATAC-seq experiment in GM12878 that was not used for QTL mapping, but was used to assess genotypic effects on the reference epigenome. ATAC-seq library preparation was performed as previously described^16^. We performed 75bp paired end sequencing in 4.4 billion sequence fragments on a HiSeq 2500 (Illumina). See Section 2 of Supplementary Note for more details.

### Sequencing data preprocessing

All sequence data sets were aligned to human genome assembly GRCh37. FASTQ files of GEUVADIS RNA-seq data^32^ (*N*=372) were downloaded from ArrayExpress (Accession E-GEUV-3), ChIP-seq data for CTCF binding^35^ (*N*=50) were downloaded from the European Nucleotide Archive (Accession ERP002168). Our ATAC-seq data and the CTCF ChIP-seq data were aligned using bwa 0.7.4^36^. RNA-seq data were aligned using Bowtie2^37^ and reads mapped to splice junctions using TopHat2^38^, using ENSEMBL human gene assembly 69 as the reference transcriptome. Following alignment, we performed peak calling in the CTCF ChIP-seq and ATAC-seq data by pooling all samples. Fragment counts of ATAC-seq, CTCF ChIP-seq and RNA-seq for each feature (a called peak or an union of exons for each gene) were normalised into FPKMs using length referred to the peak length in kilobases. Batch effects were adjusted by GC contents and principal components. See Section 3.1-3.5 of Supplementary Note for more detail.

### SNP genotype data

We downloaded VCF files from the 1000 Genomes Phase III integrated variant set from the project website (http://www.internationalgenome.org/data). For the ATAC-seq, RNA-seq and CTCF ChIP-seq samples that did not overlap with the 1000 Genomes Phase III samples, we extracted genotype data from the 1000 Genomes Phase I data or 1000 Genomes high density SNP chip data (performed on the Illumina Omni platform). We then performed whole genome imputation for the extracted genotype data by using the Beagle software^39^ (version: 23-Jul-16). See Section 3.6 in Supplementary Note for details.

### Genomic annotations

To compute ATAC-seq peak height, we pooled ATAC-seq data for the 100 samples. The peak height was defined as the highest value of the coverage depth within each peak region. Peak height quantile normalised across all peaks. The relative coverage at each variant location was calculated by the absolute coverage depth divided by the peak height inside the peak. This value was used as the variant location (VL) prior probability for both caQTL mapping and eQTL mapping. Peak distance was calculated based on the midpoint of a peak region.

We also used various external genomic annotations for comparison. The Hi-C contact map and Hi-C loops for GM12878 were obtained from Rao et al. (2004)^10^. TAD boundaries were defined as the anchor regions of a Hi-C loop. Capture Hi-C data for GM12878 was obtained from Cairns et al. (2016)^13^ and CHiCAGO^13^ software was used to extract CHi-C interactions with CHiCAGO score > 1. The H3K27ac HiChIP data for GM12878 was obtained from Mumback et al. (2017)^12^. The JuiceBox output was processed by HiCCUPS^40^ with default parameter setting to obtain the HiChIP loops. The integrated genomic segmentation annotation^27^ combining Segway^41^ and ChromHMM^42^ results was downloaded from ENCODE Project^26^ website (http://hgdownload.soe.ucsc.edu/goldenPath/hg19/encodeDCC/wgEncodeAwgSegmentation/). Each ATAC peak was labelled by one of the 7 different segmentation categories at the peak midpoint. See Section 3.7 in Supplementary Note for details.

### Pairwise hierarchical model

The pairwise hierarchical model is a product of finite mixture probabilities over all *j-k* peak pairs in 500Kb (1 ≤ *j* < *k* ≤ *J*; *J* = 277,128). The finite mixture model comprises the regional Bayes factor 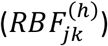 to observe chromatin accessibility *y_j_* and *y_k_* at peak *j* and *k* across 100 samples under the different interaction hypotheses *h* (Fig. 1A). The pairwise likelihood is given by

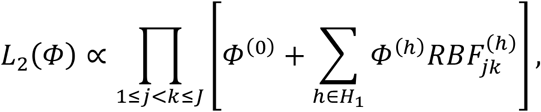

where *Φ*^(0)^ denotes the mixture probability that *j-k* peak pair is no caQTLs, *Φ*^(*h*)^ denotes the mixture probability for the alternative hypothesis *h* and *H*_1_ is the set of alternative hypotheses, so that 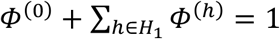. RBF is obtained from the joint regression model *p*(*y_j_, y_k_*|*h*) which comprises two independent regression models that also depend on the hypothesis *h*. For the causality hypotheses (*H*_4.1_ for the causal interaction from peak *j* to *k* and *H*_4.2_ for peak *k* to *j*), we used the two stage least square^24^ (2SLS) method to estimate the causal effect between peaks with each genetic variant in the *cis* window as the instrumental variable (Fig. 1C).

To reduce the computational complexity, we employed a two-stage optimisation of the likelihood. In the first stage we estimated hyperparameters for the variant-level and peak-level prior probabilities. We used the standard hierarchical model^23^ to learn these prior probabilities by temporarily assuming peaks are independent. In the second stage, we estimated hyperparameters in the peak-pair-level prior regarding *Φ*^(*h*)^. We used the Expectation-Maximisation algorithm to iteratively estimate hyperparameters while updating the following posterior probabilities

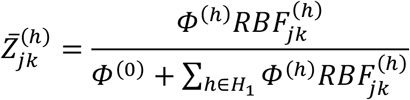

in the E-step. Because all model distributions belong to exponential family, we can utilise the penalised iteratively reweighted least square (P-IRLS) method^43^ in the M-step, which does not require calculation of the gradient and Hessian of the log likelihood. All subsequent analyses were performed based on the posterior probabilities 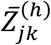 without any threshold. Note that the posterior probability of causality (PPC) is denoted by PPC_*jk*_ and PPC_*kj*_ (corresponding to 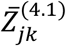 and 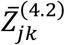, respectively) in the main text. Mathematical rationale and implementation of the pairwise hierarchical model are fully described in Section 4.1-4.5 of Supplementary Note. The software is available from GitHub (https://github.com/natsuhiko/PHM).

### Mapping multi-way interactions

Multi-way interactions were also constructed from PPC_*jk*_ and PPC_*kj*_ by finding a DAG among more than 2 peaks. We first used only confident causal interactions with PPC_*jk*_>0.5, then found the most likely parent for each peak, and finally solve the cyclic graphs by discarding an interaction with the lowest PPC_*jk*_. See Section 4.6 of Supplementary Note for details.

### Detection of lead caQTL variant

Within each cis-regulatory window (500Kb on either side of a peak), we calculated a posterior probability of each variant being the causal caQTL and obtained the maximum *a posteriori* variant as the lead variant. We used the pairwise likelihood to solve the problem that multiple caQTL variants are associated with chromatin accessibility due to strong linkage disequilibrium. The central assumption here is that variants predicted by our model to be upstream in the regulatory cascade are more likely to be causal. See Section 4.7 in Supplementary Note for details.

### Probability of Master regulator

We defined the master regulatory peak as a peak with more than one interacting downstream peak and no interacting upstream peaks. We computed the product of the following two posterior probabilities: the probability that the peak regulates at least one other peak in the *cis*-window, and the probability the peak is not regulated by any other peak within the *cis*-window, which we referred to as the probability of being master regulator (PMR). See Section 4.8 of Supplementary Note for details.

### Hierarchical model for eQTL fine-mapping

The standard hierarchical model^23^ was applied to the gEUVADIS RNA-seq data (372 European samples) with various combinations of the following five annotations: (1) inside or outside an ATAC peak (referred to as ATAC); (2) eQTL variant location, relative to an ATAC peak (referred to as variant location; VL); (3) promoter capture Hi-C contacts (CHi-C); (4) HiChIP loops from baited promoter regions (HiChIP); and (5) PMR value at each ATAC peak. The variant-level prior was learned and the posterior probability of association (PPA) was calculated for each variant in 1Mb cis-window centred at TSS. For the eQTL fine-mapping of *BLK/FAM167A* locus, we incorporated all the genomic annotations used in the caQTL mapping in conjunction with the colocalisation probability of caQTL and eQTL as the weight of the prior probability. See Section 4.9 of Supplementary Note for details.

### Colocalisation with expression QTLs

The pairwise hierarchical model can be utilised to colocalise caQTLs with other cellular QTLs, such as expression QTLs (eQTLs). The reduced model without causality hypothesis (*H*_4.1_ and *H*_4.2_) was applied to colocalise caQTL-eQTL as well as CTCF binding QTL-caQTL. We assumed a non-informative prior probability for the three different levels of hierarchy and estimated the posterior probability of pleiotropy between caQTL and eQTL/CTCF binding QTLs as the colocalisation signal. Joint colocalisation probability between eQTL and a peak pair is also calculated from the result. See Section 4.10 of Supplementary Note for details.

### Enrichment analysis with posterior probability of causal interaction

Any enrichment analysis was carried out based on PPC for all *j-k* peak pairs. We compute a 2×2 table of a binary annotation *X_jk_* (e.g., if *j-k* peak pair within TAD then *X_jk_* = 1 otherwise 0) and the existence of causality between *j-k* peak, such that

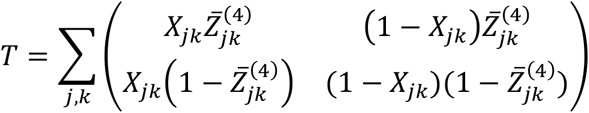

where 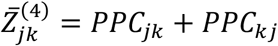 (PPC for peak *j* and *k*). We compute the odds ratio from the table *T* to perform hypothesis testing. See Section 4.11 of Supplementary Note for details.

### Knock-out of BLK-FAM167A locus (rs558245864) using CRISPR/Cas9

The lymphoblastoid cell line HG00146, which is homozygous for the reference rs558245864 allele, was nucleofected with an enhanced Cas9-2a-GFP plasmid and a guide RNA expression plasmid targeting the rs558245864 locus. Deletion clones were selected, expanded and then subjected to ATAC-seq and RNA-seq. Methods for engineering of the rs558245864 locus are described in full in the Section 5.1-5.6 of Supplementary Note.

### Differential chromatin accessibility and expression analyses

We used DESeq^44^ to perform differential chromatin accessibility and differential expression analyses. We compared the two replicates of the parental line against the four replicates of the deletion lines (two replicates for D1 and D2 heterozygous lines, respectively). See Section 5.7 of Supplementary Note for more detail.

## References

1. Pombo, A. & Dillon, N. Three-dimensional genome architecture: players and mechanisms. Nat Rev Mol Cell Biol 16, 245–57 (2015).

2. Haarhuis, J.H.I. et al. The Cohesin Release Factor WAPL Restricts Chromatin Loop Extension. Cell 169, 693–707 e14 (2017).

3. Fudenberg, G. et al. Formation of Chromosomal Domains by Loop Extrusion. Cell Rep 15, 2038–49 (2016).

4. Claussnitzer, M. et al. FTO Obesity Variant Circuitry and Adipocyte Browning in Humans. N Engl J Med 373, 895–907 (2015).

5. Smemo, S. et al. Obesity-associated variants within FTO form long-range functional connections with IRX3. Nature 507, 371–5 (2014).

6. Denker, A. & de Laat, W. The second decade of 3C technologies: detailed insights into nuclear organization. Genes Dev 30, 1357–82 (2016).

7. de Wit, E. & de Laat, W. A decade of 3C technologies: insights into nuclear organization. Genes Dev 26, 11–24 (2012).

8. Bonev, B. & Cavalli, G. Organization and function of the 3D genome. Nat Rev Genet 17, 661–678 (2016).

9. Lieberman-Aiden, E. et al. Comprehensive mapping of long-range interactions reveals folding principles of the human genome. Science 326, 289–93 (2009).

10. Rao, S.S. et al. A 3D map of the human genome at kilobase resolution reveals principles of chromatin looping. Cell 159, 1665–80 (2014).

11. Mifsud, B. et al. Mapping long-range promoter contacts in human cells with high-resolution capture Hi-C. Nat Genet 47, 598–606 (2015).

12. Mumbach, M.R. et al. Enhancer connectome in primary human cells identifies target genes of disease-associated DNA elements. Nat Genet (2017).

13. Cairns, J. et al. CHiCAGO: robust detection of DNA looping interactions in Capture Hi-C data. Genome Biol 17, 127 (2016).

14. Grubert, F. et al. Genetic Control of Chromatin States in Humans Involves Local and Distal Chromosomal Interactions. Cell 162, 1051–65 (2015).

15. Waszak, S.M. et al. Population Variation and Genetic Control of Modular Chromatin Architecture in Humans. Cell 162, 1039–50 (2015).

16. Kumasaka, N., Knights, A.J. & Gaffney, D.J. Fine-mapping cellular QTLs with RASQUAL and ATAC-seq. Nat Genet 48, 206–13 (2016).

17. Giambartolomei, C. et al. Bayesian test for colocalisation between pairs of genetic association studies using summary statistics. PLoS Genet 10, e1004383 (2014).

18. Voight, B.F. et al. Plasma HDL cholesterol and risk of myocardial infarction: a mendelian randomisation study. Lancet 380, 572–80 (2012).

19. Do, R. et al. Common variants associated with plasma triglycerides and risk for coronary artery disease. Nat Genet 45, 1345–52 (2013).

20. Day, F.R. et al. Genomic analyses identify hundreds of variants associated with age at menarche and support a role for puberty timing in cancer risk. Nat Genet 49, 834–841 (2017).

21. Day, F.R. et al. Physical and neurobehavioral determinants of reproductive onset and success. Nat Genet 48, 617–623 (2016).

22. Day, F.R. et al. Large-scale genomic analyses link reproductive aging to hypothalamic signaling, breast cancer susceptibility and BRCA1-mediated DNA repair. Nat Genet 47, 1294–1303 (2015).

23. Veyrieras, J.B. et al. High-resolution mapping of expression-QTLs yields insight into human gene regulation. PLoS Genet 4, e1000214 (2008).

24. Burgess, S. & Thompson, S.G. Mendelian randomization: methods for using genetic variants in causal estimation. in Chapman & Hall/CRC interdisciplinary statics series 1 online resource (CRC Press, Taylor & Francis Group,, Boca Raton, FL, 2015).

25. Ignatiadis, N., Klaus, B., Zaugg, J.B. & Huber, W. Data-driven hypothesis weighting increases detection power in genome-scale multiple testing. Nat Methods 13, 577–80 (2016).

26. Consortium, E.P. An integrated encyclopedia of DNA elements in the human genome. Nature 489, 57–74 (2012).

27. Hoffman, M.M. et al. Integrative annotation of chromatin elements from ENCODE data. Nucleic Acids Res 41, 827–41 (2013).

28. Tewhey, R. et al. Direct Identification of Hundreds of Expression-Modulating Variants using a Multiplexed Reporter Assay. Cell 165, 1519–1529 (2016).

29. Guthridge, J.M. et al. Two functional lupus-associated BLK promoter variants control cell-type- and developmental-stage-specific transcription. Am J Hum Genet 94, 586–98 (2014).

30. Consortium, G.T. et al. Genetic effects on gene expression across human tissues. Nature 550, 204–213 (2017).

31. Battle, A. et al. Characterizing the genetic basis of transcriptome diversity through RNA-sequencing of 922 individuals. Genome Res 24, 14–24 (2014).

32. Lappalainen, T. et al. Transcriptome and genome sequencing uncovers functional variation in humans. Nature 501, 506–11 (2013).

33. Shin, H.Y. et al. Hierarchy within the mammary STAT5-driven Wap superenhancer. Nat Genet 48, 904–911 (2016).

34. Chen, L. et al. Genetic Drivers of Epigenetic and Transcriptional Variation in Human Immune Cells. Cell 167, 1398–1414 e24 (2016).

35. Ding, Z. et al. Quantitative genetics of CTCF binding reveal local sequence effects and different modes of X-chromosome association. PLoS Genet 10, e1004798 (2014).

36. Li, H. & Durbin, R. Fast and accurate short read alignment with Burrows-Wheeler transform. Bioinformatics 25, 1754–60 (2009).

37. Langmead, B. & Salzberg, S.L. Fast gapped-read alignment with Bowtie 2. Nat Methods 9, 357–9 (2012).

38. Kim, D. et al. TopHat2: accurate alignment of transcriptomes in the presence of insertions, deletions and gene fusions. Genome Biol 14, R36 (2013).

39. Browning, B.L. & Browning, S.R. Genotype Imputation with Millions of Reference Samples. Am J Hum Genet 98, 116–26 (2016).

40. Durand, N.C. et al. Juicer Provides a One-Click System for Analyzing Loop-Resolution Hi-C Experiments. Cell Syst 3, 95–8 (2016).

41. Hoffman, M.M. et al. Unsupervised pattern discovery in human chromatin structure through genomic segmentation. Nat Methods 9, 473–6 (2012).

42. Ernst, J. & Kellis, M. ChromHMM: automating chromatin-state discovery and characterization. Nat Methods 9, 215–6 (2012).

43. Wood, S.N. Generalized additive models: an introduction with R, xvii, 391 p. (Chapman & Hall/CRC, Boca Raton, Fla.; London, 2006).

44. Anders, S. & Huber, W. Differential expression analysis for sequence count data. Genome Biol 11, R106 (2010).

